# PINT: Pathway-pathway interactions for predicting interpretable clinical outcomes from gene expression

**DOI:** 10.64898/2026.08.11.744286

**Authors:** Sai Phani Parsa, Beomsu Baek, Euiseong Ko, Sai Chandra Kosaraju, Mingon Kang

## Abstract

**Motivation:** Disease mechanisms emerge from the coordinated activity of multiple biological pathways, rather than from individual pathways acting in isolation. Existing pathway-based deep learning models, however, treat pathways as independent entities, aggregating their representations through fully connected layers that disregard inter-pathway relationships. This architectural limitation overlooks an important dimension of disease biology, potentially constraining both predictive performance and the capacity to generate biologically meaningful interpretations.

**Results:** We introduce a pathway-based attentive interpretability model, named PINT, that models interactions among pathways through a self-attention mechanism from gene expression data. An attention-based pooling layer further identifies patient-specific pathway contributions to the final prediction. Evaluation across five TCGA cancer datasets demonstrated that PINT consistently outperformed benchmark models in survival analysis. More importantly, PINT identifies pathways significantly associated with survival as well as reveals biologically meaningful interactions among pathways. In the BRCA dataset, PINT identified significant pathways, pathway-pathway interactions, and gene-level contributions within pathways for individual patients, most of which were supported by existing literature. Specifically, the RAS signaling pathway emerged as significantly associated with patient survival, and the learned interaction scores recovered known relationships between RAS signaling and several regulatory pathways, including cAMP, TNF, and Rap1 signaling.

**Availability and implementation:** The source code and data are available at https://github.com/datax-lab/PINT.

## 1. Introduction

Biological pathways are coordinated molecular processes that regulate cellular functions to define the functional state of cells. These pathways operate in an interconnected manner; components of one pathway influence elements of another through complex interactions between pathways (Creixell et al., 2015). The equilibrium among these interacting pathways helps maintain cellular homeostasis, while disturbances in their relationships can disrupt normal cellular activity and contribute to the onset and progression of disease (Liu et al., 2024; Fang et al., 2019). Such disturbances are common in cancer, where interactions among signaling and regulatory pathways have been linked to tumor development, metastasis, and therapeutic response (Sever and Brugge, 2015). Thus, coordinated interactions between pathways may better describe the molecular complexity of these conditions than abnormalities in any single pathway alone (Zheng et al., 2020).

Most of the existing pathway-based deep learning models have represented each pathway as an isolated unit. They typically incorporate biological pathway knowledge into their architectures to constrain high-dimensional gene expression data into biologically meaningful representations by explicitly mapping genes to their corresponding pathways. Within these frameworks, the models construct multiple pathway-level embeddings, each computed from the subset of genes that belong to a given pathway, and combine these embeddings to predict clinical outcomes (Hao and Kim, 2018; Hao et al., 2018; Ko et al., 2024, 2026). Pathway-specific encoders learn independent, pathway-tailored representations based on the expression profiles of the genes assigned to each pathway (Park et al., 2022). Hierarchical architectures further organize genes and pathways into multi-tiered structures that mirror biological hierarchies, allowing information to be transmitted across progressively more abstract levels (Elmarakeby et al., 2021). However, embedding an explicit representation of these inter-pathway dependencies has the potential to improve predictive performance and, crucially, to yield interpretable explanations of which specific pathway interactions underlie a given clinical outcome.

In this study, we introduce a pathway-based attentive interpretability framework, named PINT, that explicitly models interactions among biological pathways for predicting clinical outcomes. PINT employs an attention mechanism to capture pathway–pathway interactions from gene expression profiles, enabling the model to identify the most relevant pathway interactions for a specific prediction. This architecture offers interpretable insights into how interacting pathways influence clinical outcomes, advancing beyond the single-pathway interpretations provided by current methods.

## 2. Methods

The PINT architecture consists of four components: (1) a pathway-specific representation layer, (2) a pathway-pathway interaction layer, (3) a sample-level representation layer, and (4) an output layer (Fig. 1). First, the pathway-specific representation layer generates pathway-level embeddings from gene expression profiles by leveraging prior knowledge of gene–pathway associations. In this layer, a dedicated encoder is assigned to each pathway, which learns its representation based on the expression patterns of the genes linked to that pathway. Second, the pathway-pathway interaction layer captures coordinated activity across pathways by using a multi-head self-attention mechanism over the learned pathway representations to model their mutual interactions. Third, pathway embeddings are combined via an attention-driven pooling strategy at the sample representation level, allowing the model to identify which pathways have the most substantial impact on a given prediction. Fourth, the output layer forwards the derived sample-level representation into fully connected, task-tailored layers, which then generate either prognostic indices or survival classification probabilities, contingent upon the specific downstream objective.

**Figure 1.**
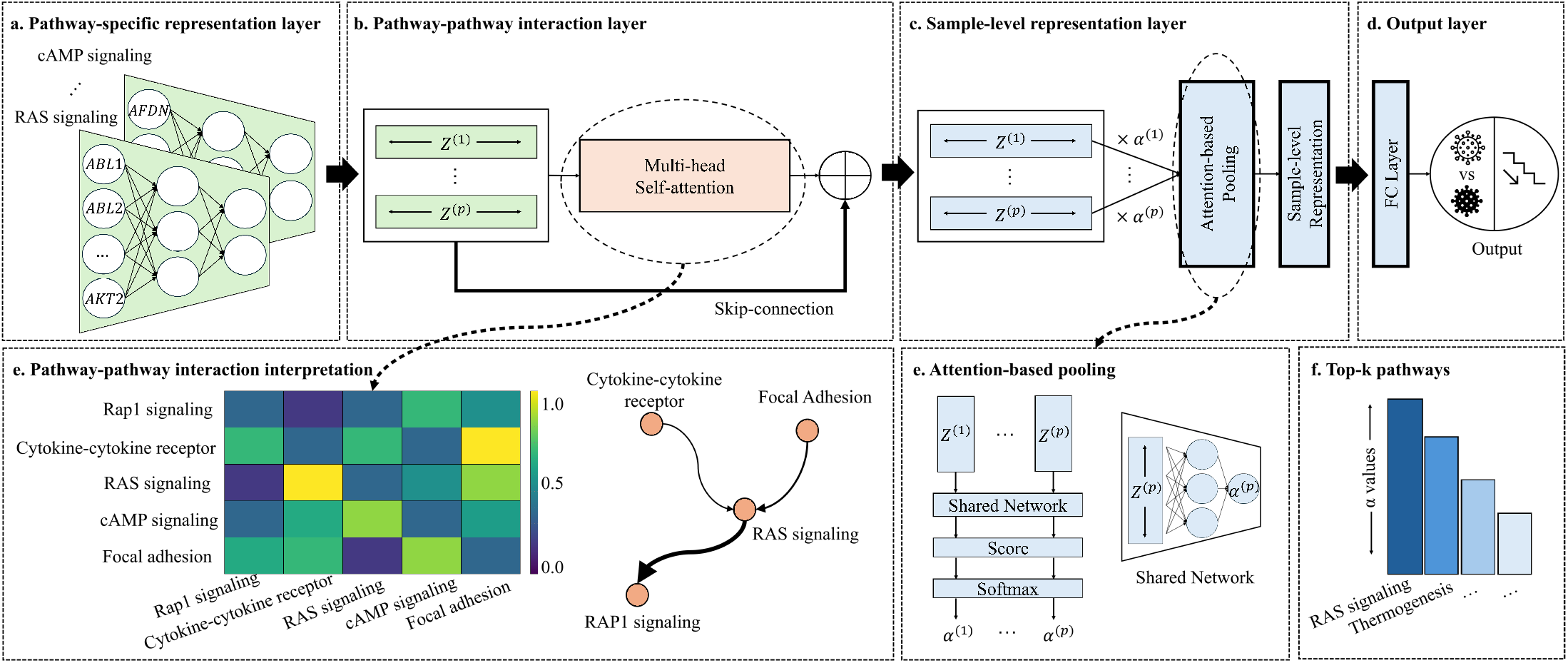
Overview of the proposed PINT architecture. (a) Gene expression features are encoded using pathway-specific encoders to produce pathway embeddings. (b) Multi-head self-attention models pathway-pathway interactions. (c) Attention-based pooling aggregates pathway embeddings into a sample-level representation. (d) Task-specific output layers (e.g., survival analysis or classification) generate the final prediction. (e–f) Interpretability through pathway importance scores and pathway–pathway interaction maps derived from attention weights.

For each sample, the pathway-specific representation layer constructs a pathway embedding matrix **Z**_*i*_ ∈ *ℜ*^*p*×*d*^ from the gene expression matrix **X** ∈ *ℜ*^*n*×*m*^, where *n* is the number of samples and *m* is the number of genes. Let *P*_*j*_ denote the *j*^th^ biological pathway for *j* = 1, …, *p*, with *p* indicating the total number of pathways. For the *i*^th^ sample, we extract from **X** the expression values of the genes assigned to pathway *P*_*j*_, resulting in a gene-level feature vector 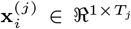, where *T*_*j*_ is the number of genes associated with pathway *P*_*j*_ . This vector is then transformed into a fixed-length representation by a pathway-specific encoder *f*_*j*_ :

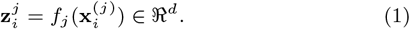

Aggregating the embeddings over all *p* pathways produces the matrix:

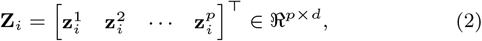

where each row encodes the representation of a single pathway, and *d* denotes the dimensionality of the embedding space.

The pathway–pathway interaction layer takes the pathway embedding matrix **Z**_*i*_ ∈ *ℜ*^*p*×*d*^ from the previous stage, as input, and models dependencies among pathways using a multi-head self-attention mechanism. For each attention head *h* = 1, …, *H*, we form the query, key, and value matrices as

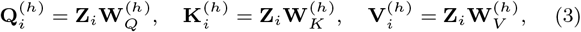

where 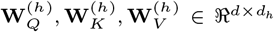 are learnable linear projection matrices, and *d*_*h*_ = *d/H* denotes the dimensionality per head. The pairwise interactions among pathways are encoded through the attention matrix

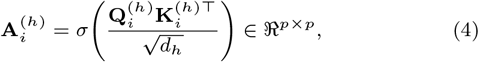

where *σ*(·) is a row-wise normalization operator that determines how attention mass is distributed over pathway pairs. In our implementation, we instantiate *σ* as the softmax function. Consequently, each entry in 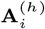 measures the strength of interaction between a particular pair of pathways in the *i*^th^ sample. The attention output for head *h* is computed as 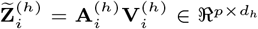. The outputs from all heads are then concatenated and passed through a linear transformation:

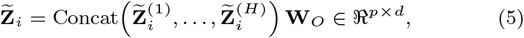

where 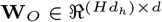 is a trainable output projection. Finally, we apply a residual connection to revise the pathway embeddings:

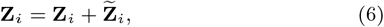

so that the updated **Z**_*i*_ retains the original pathway-specific information while being enriched with signals capturing inter-pathway interactions.

The sample-level representation layer condenses the pathway embedding matrix **Z**_*i*_ for sample *i* into a single embedding vector **s**_*i*_ via an attention-based pooling mechanism. A lightweight attention module *g*(·) assigns a relevance score to each pathway, which is subsequently transformed into normalized attention coefficients:

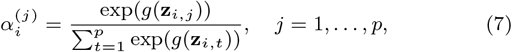

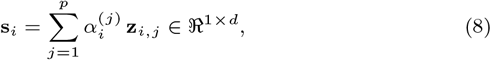

where 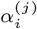 denotes the normalized attention weight assigned to pathway *P*_*j*_ in sample *i*, reflecting its relative contribution to the resulting sample-level embedding.

The output module then projects **s**_*i*_ through one or more fully connected layers, followed by a task-specific prediction head. With this architecture, the PINT framework can be adapted to a variety of downstream tasks, including survival analysis, survival risk stratification, and recurrence prediction. In the survival analysis setting considered in this work, the output layer generates a prognostic index, which is optimized by minimizing the negative Cox partial log-likelihood loss *ℒ*_Cox_.

To encourage the model to focus on a small set of highly informative pathways, instead of spreading attention across many, we add an entropy-based regularization term to the attention weights used in pooling. For a given sample *i*, we define the normalized entropy as

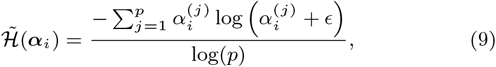

where ϵ is a small constant included for numerical stability, and the factor log(*p*) normalizes the entropy by the number of pathways, yielding scale-invariant values. The overall training objective augments the survival loss with the average entropy penalty across a mini-batch of size *B*:

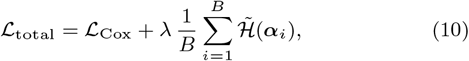

where *λ* determines how strongly the regularization term is applied. By reducing this entropy term, the model is encouraged to produce attention distributions that more clearly emphasize a limited set of pathways, thereby increasing pathway-level sparsity and improving the interpretability of the resulting sample embeddings.

### 2.1. Model Interpretation

PINT achieves interpretability through the attention mechanisms in both the pathway–pathway interaction layer and the sample-level representation layer. In the sample-level representation layer, an importance weight 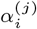 is assigned to each pathway for each sample, indicating the extent to which that pathway contributes to the final prognostic prediction for that sample. In parallel, the pathway–pathway interaction layer outputs an attention matrix 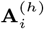, whose entries measure the interaction strength between each pair of pathways for a given sample. Together, these two attention modules enable interpretation at the level of individual pathways and their pairwise interactions.

Pathway relevance is assessed by ranking pathways according to their attention weights from the sample-level representation layer and examining how consistently they are selected across the dataset. For each sample *i*, the index set of the top *K* highest-weighted pathways is defined as 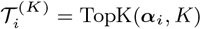, where *K* is a user-specified number of top-ranked pathways. The number of times each pathway *j* is selected across *n* samples is then given by

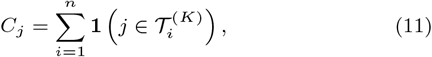

where **1**(·) denotes the indicator function, and the empirical selection rate is *R*_*j*_ = *C*_*j*_ */n*. To test whether a pathway is selected more often than expected by random chance, a null model of uniform random selection is adopted. Under this null, each sample corresponds to a Bernoulli trial with success probability *p*_0_ = *K/p*, and the total selection count follows *C*_*j*_ ∼ Binomial(*n, p*_0_). Pathways with selection frequencies that remain significant after FDR-BH multiple testing correction are deemed robustly influential.

Inter-pathway interaction scores are derived from the attention matrices output by the pathway-pathway interaction layer. For each sample, the head-specific attention matrices **A**^(*h*)^ ∈ *ℜ*^*p*×*p*^ are combined across heads as

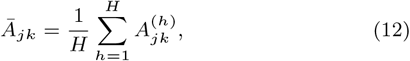

yielding a sample-specific interaction matrix **Ā** ∈ *ℜ*^*p*×*p*^, where *Ā*^*jk*^ represents the degree to which pathway *k* contributes to the representation of pathway *j*. To isolate cross-pathway effects, all diagonal entries are set to zero.

To highlight interactions involving pathways that are most relevant for prediction, a relevance-weighted interaction score fuses the two attention sources. For a given sample, let *α*^(*j*)^ be the attention weight assigned to pathway *j* by the sample-level representation layer. The relevance-weighted directional interaction score is then defined as

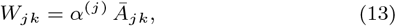

where *W*_*jk*_ encodes how strongly pathway *j* attends to pathway *k*, scaled by the predictive importance of pathway *j*. For each sample, this yields an interaction matrix **W** ∈ *ℜ*^*p*×*p*^ that summarizes pathway dependencies while incorporating their relevance to the predictive task. Outgoing and incoming interaction profiles for any pathway can be extracted from the corresponding row and column of **W**, respectively, allowing the identification of pathways that exert strong influence on, or are strongly influenced by, a pathway of interest.

## 3. Results

### 3.1. PINT improves predictive performance compared with existing benchmark models

We benchmarked PINT against six established survival analysis methods: the penalized Cox proportional hazards model Cox-EN (Simon et al., 2011); the tree-ensemble method Random Survival Forests (RSF) (Ishwaran et al., 2008); the deep learning models DeepSurv (Katzman et al., 2018) and AESurv (Shen et al., 2024); and the pathway-informed approaches CoxPasNet (Hao et al., 2018) and DeepHisCoM (Park et al., 2022). Together, these benchmark models encompass classical statistical modeling, machine learning, deep neural networks, and pathway-centric frameworks, thereby enabling a comprehensive comparison across distinct methodological families. Performance comparison experiments were performed in five cohorts of TCGA cancer: lower-grade brain glioma (LGG), breast invasive carcinoma (BRCA), liver hepatocellular carcinoma (LIHC), kidney renal clear cell carcinoma (KIRC) and lung adenocarcinoma (LUAD). All datasets were retrieved from the cBioPortal for Cancer Genomics (cBioPortal for Cancer Genomics, 2012) representing heterogeneous malignancies with differing cohort sizes (Table 1; data preprocessing procedures are detailed in the Supplementary Note S1). To encode prior biological knowledge, we incorporated human pathway annotations from the KEGG database (Kyoto Encyclopedia of Genes and Genomes, 2025), retaining 231 pathways after applying filters based on biological interpretability and gene set cardinality (see Supplementary Note S2 for the curation criteria).

**Table 1.**
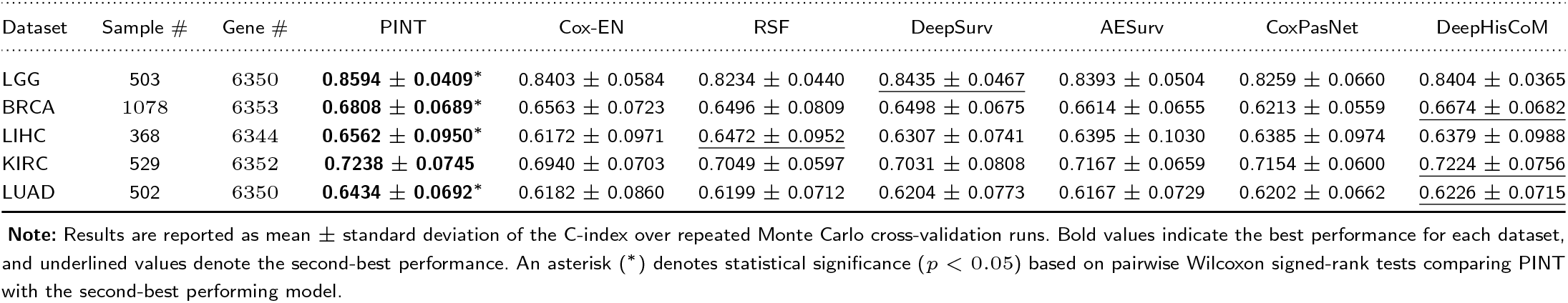
Survival analysis performance across TCGA datasets measured by C-index (mean *±* std).

Within each cohort, the data were split into training (80%), validation (10%), and test (10%) sets, while preserving the proportion of censored and uncensored observations. Normalization parameters were estimated exclusively from the training data and then applied to the validation and test sets to prevent information leakage. We computed concordance index (C-index) for the performance comparison. We optimized hyperparameters that maximized C-index on the validation set for PINT and all benchmark models. For PINT, the hyperparameters included the number of attention heads *H*, the embedding dimension *d*, and the regularization coefficient *λ* (complete search grids and chosen settings are reported in Supplementary Note S3). Each experimental setup was repeated 10 times with independent random splits of the data, and differences in performance were evaluated for statistical significance using the Wilcoxon signed-rank test.

PINT consistently achieved the highest mean C-index in all five TCGA datasets, with values of 0.8594 ± 0.0409 in LGG, 0.6808 ± 0.0689 in BRCA, 0.6562 ± 0.0950 in LIHC, 0.7281 ± 0.0745 in KIRC, and 0.6434 ± 0.0692 in LUAD (Table 1) The improvement over the benchmark models was up to 3.34%, with consistent gains across all the datasets. These results were further supported by the Wilcoxon rank-sum test, which indicated statistically significant improvements in LGG, BRCA, LIHC, and LUAD. The consistency of these gains across cancer types with varying sample sizes and molecular characteristics suggests that PINT generalizes effectively across heterogeneous settings.

To further assess generalization, we applied the trained BRCA models to the independent METABRIC-BRCA dataset, which contains gene expression profiles and clinical survival information for breast cancer patients. For each model, the best-performing checkpoint from the 10 repeats was selected based on the validation C-index and applied to the external dataset. PINT achieved the highest C-index (0.5852 ± 0.0319) compared with DeepHisCoM (0.5781 ± 0.0351), DeepSurv (0.5749 ± 0.0084), AESurv (0.5721 ± 0.0399), CoxPasNet (0.5703 ± 0.0339), RSF (0.5616 ± 0.0262), and Cox-EN (0.5605 ± 0.0119) (Table 2) This trend is consistent with results from the TCGA-BRCA dataset, indicating that PINT’s performance advantage may extend beyond the training distribution.

**Table 2.**
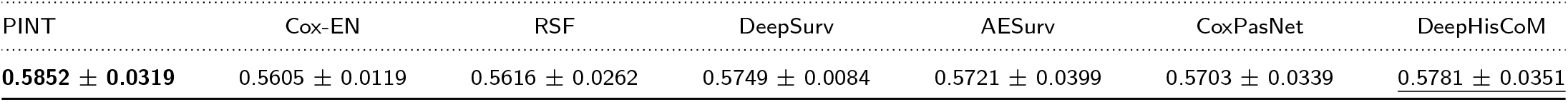
Generalization performance on the METABRIC-BRCA dataset measured by C-index (mean *±* std).

| PINT | Cox-EN | RSF | DeepSurv | AESurv | CoxPasNet | DeepHisCoM |
| --- | --- | --- | --- | --- | --- | --- |
| **0.5852 ± 0.0319** | 0.5605 ± 0.0119 | 0.5616 ± 0.0262 | 0.5749 ± 0.0084 | 0.5721 ± 0.0399 | 0.5703 ± 0.0339 | <u>0.5781 ± 0.0351</u> |

### 3.2. Ablation study

To quantify the contribution of each architectural element in PINT, we conducted an ablation study on the BRCA dataset, comparing the complete model to two reduced variants. In the first variant, the multi-head self-attention module was removed, and pathway embeddings were fed directly into the attention-based pooling layer, thereby omitting explicit modeling of pathway–pathway dependencies. In the second variant, the attention-based pooling layer was replaced with simple mean pooling over the self-attention layer’s outputs to derive sample-level representations. For all model configurations, hyperparameters were tuned using Optuna (Akiba et al., 2019).

The full PINT architecture attained a C-index of 0.6808 ±0.0689, exceeding the performance of the variant without multi-head self-attention (0.6271 ± 0.1160) by 8.5% and that of the variant without attention-based pooling (0.6532 ± 0.0512) by 4.2% (Table 3) The performance gain relative to the no–self-attention variant was statistically significant (*p* < 0.05), whereas the improvement over the no–attention-based-pooling variant was borderline significant (*p* = 0.0527). Collectively, these findings indicate that both components contribute in a complementary fashion, with the multi-head self-attention module providing the dominant benefit by capturing inter-pathway relationships that are not reflected in isolated pathway embeddings.

**Table 3.**
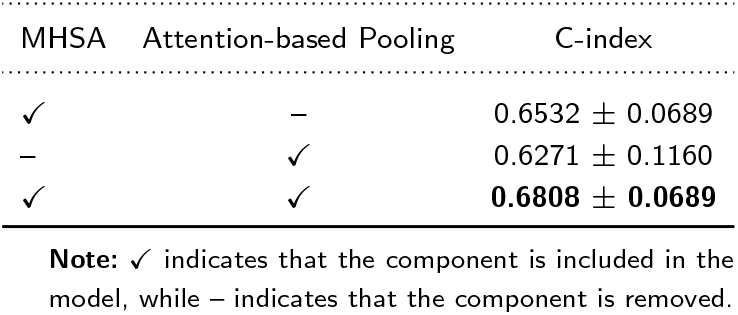
Ablation study evaluating the contribution of the multi-head self-attention and attention-based pooling modules on the BRCA dataset. Performance is reported as C-index (mean ± std).

| MHSA | Attention-based Pooling | C-index |
| --- | --- | --- |
| ✓ | – | 0.6532 ± 0.0689 |
| – | ✓ | 0.6271 ± 0.1160 |
| ✓ | ✓ | **0.6808 ± 0.0689** |

**Note:** ✓ indicates that the component is included in the model, while – indicates that the component is removed.

### 3.3. PINT identifies statistically significant pathways associated with breast cancer

We identified top-10 pathways contributing to patient survival in breast cancer by interpreting PINT. Attention weights 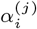 from the sample level representation layer were collected for all test samples across experimental runs, and when a sample appeared in multiple test sets, its attention weights were averaged to obtain a single representative pathway score. Pathways were then ranked for each sample in descending order by attention weight, and we computed how often each pathway appeared among the top *K* positions, with *K* = 10 chosen to balance emphasis on the most influential pathways with sufficient occurrences for reliable statistical testing.

Using the binomial test with FDR BH correction, we identified 31 pathways that were selected more frequently than expected by chance. The 10 most frequently selected pathways are shown in Table 4 and Fig. 2. To assess whether these pathway importance scores have prognostic value, we conducted univariate Cox proportional hazards analysis using the *z* score normalized attention weights as continuous covariates. We fitted a separate Cox model for each pathway, estimating the hazard ratio (HR) and 95 percent confidence interval, and assessed significance using the Wald test (Davidson-Pilon, 2019). After FDR BH correction (*q* < 0.10), two pathways showed significant associations with survival: the RAS signaling pathway and Endocytosis, both indicating increased risk (*HR* > 1), as presented in Fig. 2b.

**Table 4.** Top-10 statistically significant pathways identified by PINT.

| Pathway | Supporting Evidence (PMID) |
| --- | --- |
| Natural killer cell-mediated cytotoxicity | 35116714 |
| Thermogenesis | 34071012 |
| Endocytosis | 38769215 |
| Apoptosis | 38201650 |
| Pentose and glucuronate interconversions | – |
| RAS signaling pathway | 31781501, 40500484 |
| Longevity regulating pathway | – |
| Phospholipase D signaling pathway | 33983572 |
| Aminoacyl tRNA biosynthesis | – |
| JAK-STAT signaling pathway | 40975142, 35116714 |

**Note:** PMID denotes the PubMed identifier of studies reporting evidence linking the corresponding pathway to breast cancer progression or prognosis. A dash (–) indicates that direct supporting literature was not identified in our survey.

**Figure 2.**
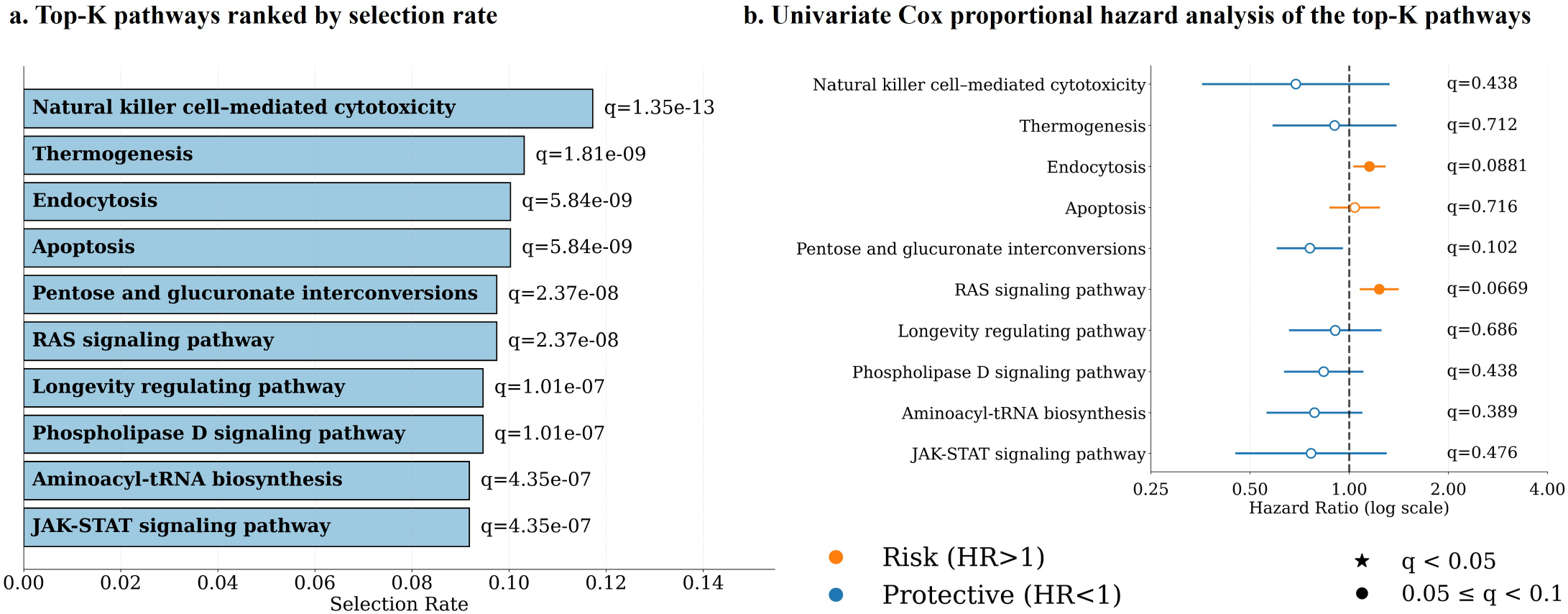
Top-*K* pathway interpretation results on the TCGA-BRCA dataset. (a) Top-10 pathways ranked by selection rate, where selection rate denotes the fraction of patients for which a pathway appeared among the top-*K* pathways based on attention-based pooling scores. The reported *q*-values are FDR-adjusted. (b) Univariate Cox proportional hazards analysis of the same top-10 pathways using pathway attention scores as continuous covariates. Points indicate hazard ratios, horizontal bars denote 95% confidence intervals, and colors distinguish risk-associated pathways (HR *>* 1) from protective pathways (HR *<* 1). The reported *q*-values are FDR-adjusted, and two pathways remained significant at *q <* 0.10.

Of these, the RAS signaling pathway is particularly relevant to breast cancer. Dysregulated RAS signaling has repeatedly been associated with tumor progression, metastatic spread, and resistance to therapy in breast cancer (Wu et al., 2025; Galiè, 2019), and its identification as the top survival-associated pathway by Path AIM supports the biological plausibility of the model representations. Endocytic activity has likewise been linked to aggressive disease behavior, with elevated expression of MAL2, a regulator of endocytic trafficking, independently predicting shorter survival in TCGA BRCA and triple negative breast cancer cohorts (Borowczak et al., 2024). Several other pathways highlighted by Path AIM, including Thermogenesis, JAK-STAT signaling, natural killer cell-mediated cytotoxicity, Apoptosis, and Phospholipase D signaling, have also been reported to affect breast cancer progression, tumor immune interactions, or clinical outcomes in previous studies (Gandhi et al., 2021; Feng et al., 2025; Wang et al., 2021; Pandey et al., 2024; Lee et al., 2021).

### 3.4. Pathway-pathway interactions

We futher analyzed pathway–pathway interactions, focusing on the RAS signaling pathway, which was identified as significantly associated with breast cancer survival. As a central driver of tumor growth, metastatic spread, and treatment resistance in breast cancer, the RAS signaling pathway represents an informative hub for examining how pathways interact to shape clinical outcomes.

Analysis of interaction scores from the pathway-pathway interaction layer revealed several biologically coherent interactions with the RAS signaling pathway (Fig. 3). Among the strongest interactions, Rap1 signaling emerged as a prominent partner. This is consistent with the KEGG pathway map for RAS signaling, which positions Rap1 as a signaling node linking RAS-mediated signaling to the dedicated Rap1 signaling pathway (Kyoto Encyclopedia of Genes and Genomes, 2025), confirming that PINT recovers known inter-pathway relationships.

**Figure 3.**
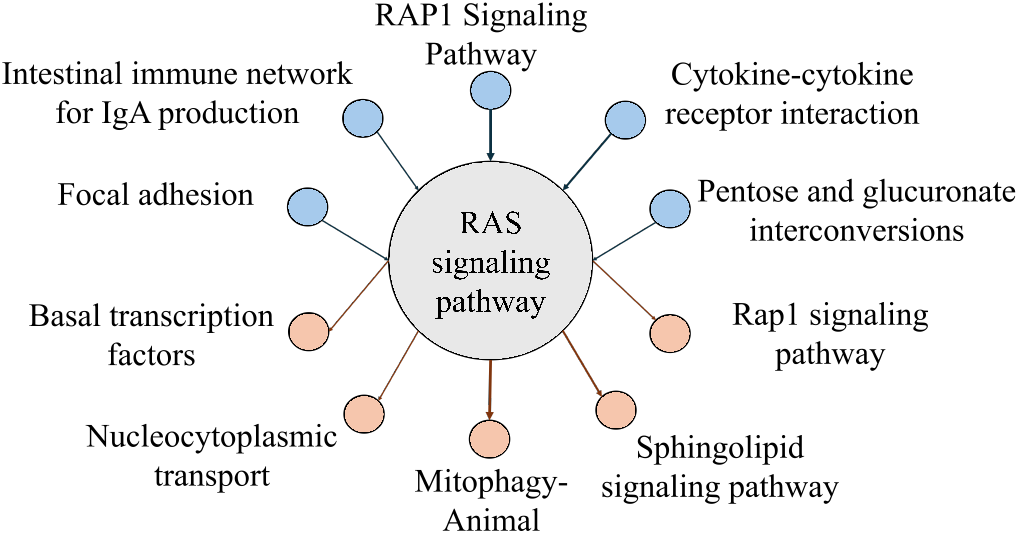
Pathway–pathway interactions involving the RAS signaling pathway. The top five pathways influencing the RAS signaling pathway (blue) and the top five pathways influenced by RAS signaling (orange) identified by the model are shown. Nodes represent biological pathways and directed edges indicate inferred pathway–pathway interactions, with the RAS signaling pathway highlighted at the center.

Inflammatory cytokine and focal adhesion pathways were also identified as strongly interacting with RAS signaling. TNF*α* and IL-1*β*, components of inflammatory cytokine signaling, have been reported to activate RAS signaling and cooperate with RAS to promote tumor progression (Leibovich-Rivkin et al., 2014). Focal adhesion kinase activity has similarly been correlated with RAS-associated tumor growth (Sulzmaier et al., 2014).

PINT also identified interactions between RAS signaling and TNF signaling, as well as cytoskeletal regulation pathways. These relationships have been described in prior studies: RAS and TNF signaling interact through cAMP signaling networks (Stork and Schmitt, 2002; Cordero et al., 2010), and RAS signaling has been linked to cytoskeletal regulation through Rho-family signaling pathways that control actin dynamics (Soriano et al., 2021).

Collectively, the pathway interactions learned by PINT align with established biological knowledge, providing evidence that the model captures meaningful inter-pathway dependencies rather than spurious associations.

## 4. Discussion

We introduce PINT that models interactions among biological pathways for clinical outcome prediction. PINT learns pathway-level embeddings from gene expression data, employs a multi-head self-attention module to capture interactions between pathways, and uses an attention-based pooling layer to quantify patient-specific pathway contributions.

PINT consistently surpassed traditional statistical models, standard machine learning methods, and other pathway-based approaches across five TCGA cancer cohorts. The ablation analysis demonstrated that the self-attention component—tasked with modeling inter-pathway dependencies—was the primary driver of this performance gain, indicating that coordinated pathway activity carries predictive information beyond that provided by individual pathway features alone.

PINT provides interpretability at two complementary scales. At the pathway scale, it identified the RAS signaling pathway as one of the most impactful pathways in breast cancer, and Cox regression further confirmed its strong correlation with patient survival. At the interaction scale, PINT revealed dependencies between RAS signaling and multiple regulatory pathways, including cAMP, TNF, and Rap1 signaling. These inferred connections are consistent with known biological relationships, and the fact that they arise purely from learned attention weights—without any explicit prior encoding—supports the view that PINT learns biologically meaningful structure across pathways.

The pathway–pathway relationships characterized by PINT provide biomedical researchers with a data-driven framework for examining how coordinated pathway dynamics influence clinical outcomes, thereby supporting the discovery of combinatorial therapeutic targets that modulate interacting pathways instead of focusing on single pathways in isolation. Moreover, integrating multi-omics modalities could enable the model to learn regulatory relationships across different biological layers, potentially yielding deeper insight into disease mechanisms and enhancing patient stratification.

## 5. Funding

This research was supported by the DOE Office of Science, Office of Biological and Environmental Research (BER) (Grant#: DE-SC0025298), the National Science Foundation Major Research Instrumentation (NSF MRI) (Grant#: 2117941), and Cal Poly Pomona, Start-up fund (S.C.K).

## Notes

### Competing Interest Statement

The authors have declared no competing interest.

