## Supplementary for "PINT: Pathway-pathway interactions for predicting interpretable clinical outcomes from gene expression"

### Supplementary information

### Supplementary Notes

#### Supplementary Note 1: TCGA data collection and preprocessing

We evaluated Path-AIM using five cancer datasets from The Cancer Genome Atlas (TCGA): brain lower-grade glioma (LGG), breast invasive carcinoma (BRCA), liver hepatocellular carcinoma (LIHC), kidney renal clear cell carcinoma (KIRC), and lung adenocarcinoma (LUAD). These datasets were obtained from cBioPortal for Cancer Genomics (cBioPortal for Cancer Genomics, 2012), specifically from the TCGA Firehose Legacy cohorts. For each cancer type, mRNA gene expression profiles were collected along with overall survival time and event status.

Gene expression data were preprocessed as follows. Hugo gene symbols were used as feature identifiers, and the expression matrix was transposed so that rows corresponded to samples and columns to genes. Genes with missing symbols or missing expression values across all samples were removed. Duplicate gene symbols were resolved by averaging expression values across columns corresponding to the same gene, resulting in a single expression value per gene.

Clinical data were processed by retaining the sample ID, overall survival time, and survival status. Samples were restricted to primary tumors of the target cancer type, and samples with vial identifier B were excluded to avoid duplicate measurements. Records with missing survival information or negative survival times were removed. Survival status was converted to a binary variable (0 = censored, 1 = deceased), and survival times were stored as numeric values. The processed gene expression matrix and clinical data were merged using sample IDs to obtain the final dataset for each cancer type.

#### Supplementary Note 2: KEGG pathway data

Pathway annotations were obtained from the Kyoto Encyclopedia of Genes and Genomes (Kyoto Encyclopedia of Genes and Genomes, 2025), which curates' molecular interaction and reaction networks. From an initial set of 365 human (Homo sapiens) pathways, disease-specific pathways were removed to reduce bias toward disease-specific annotations and to focus on general biological processes, resulting in 257 pathways. This filtering enables the model to capture pathway activity through fundamental signaling and metabolic processes that reflect broader cellular responses. Pathways were further filtered by gene set size, removing those with fewer than 15 or more than 300 genes, yielding a final set of 231 pathways for downstream analysis.

#### Supplementary Note 3: Hyperparameter tuning and Benchmark implementations

All models were tuned using Optuna (Akiba et al., 2019) with the validation C-index as the selection criterion, ensuring a fair comparison across methods.

The Path-AIM search space covered the pathway encoder, self-attention module, and classification head. The encoder used two fully connected layers with hidden dimensions drawn from {32, 16} and {32, 16, 8}. The self-attention module used a single layer with 1, 2, or 4 attention heads, and the classification head dimension was selected from $\{32, 16, 8\}$. Dropout rates were tuned separately for each component, ranging from 0.00 to 0.30 per layer. Training used the AdamW optimizer with batch sizes of 256 or 512, learning rates between $1e-5$ $and$ $9e-5$, and weight decay from {1e-4, 3e-4, 1e-3, 3e-3, 1e-2}. A 30-epoch warm-up was followed by cosine annealing to 1e-7. Models were trained for at least 200 epochs with early stopping based on validation C-index (patience of 20). The entropy regularization coefficient $\lambda$ was selected from {1e-4, 3e-4, 1e-3, 3e-3, 1e-2}.

The Cox-EN baseline used the elastic-net penalized Cox model from scikit-survival (Pölsterl, 2020), tuning the mixing parameter and selecting the regularization strength from the regularization path. RSF was also implemented via scikit-survival (Ishwaran et al., 2008, Pölsterl, 2020), with the number of estimators, tree depth, split and leaf constraints, and feature sampling tuned on the validation set.

DeepSurv (Katzman et al., 2018) and AESurv (Shen et al., 2024) both used two hidden-layer architectures with layer sizes, dropout, learning rate, weight decay, and scheduling parameters optimized on the validation set.

CoxPasNet (Hao et al., 2018) and DeepHisCoM (Park et al., 2022) were tuned following the recommended settings from their respective repositories, optimizing learning rate, regularization, and architecture parameters on the validation set.
